# Plant quarantine alarm: as much as 20 new alien insect pest species including *Drosophila suzukii* appeared in the Caucasus in the last seven years

**DOI:** 10.1101/264127

**Authors:** Andrzej O. Bieńkowski, Marina J. Orlova-Bienkowskaja, Natalia N Karpun

## Abstract

In 2011-2017 an unusually high number of invasive pests new to European Russia were detected for the first time in Sochi on the Black Sea coast of the Caucasus. We present the first reports of two pests new for the Caucasus and European Russia found in 2017: *Drosophila suzukii* (a pest of fruit, included to EPPO A2 list) and *Otiorhynchus armadillo* (a pest of agricultural and ornamental plants). Other recently established insects: a polyphagous pest *Halyomorpha halys* (first record in 2014); pests of palm trees included to EPPO A2 list: *Paysandisia archon* (2014) and *Rhynchophorus ferrugineus* (2012); a pest of Solanaceae: *Epitrix hirtipennis* (2013); a pest of ornamental flowers: *Luperomorpha xanthodera* (2016); a pest of soybeans: *Medythia nigrobilineata* (2016); a pest of wine production: *Harmonia axyridis* (2012); a pest of strawberry: *Stelidota geminata* (2013); pests of *Eucalyptus: Ophelimus maskelli* (2011), *Glycaspis bremblecomblei* (2014), *Leptocybe invasa* (2014); a pest of Cupressaceae: *Lamprodila festiva* (2013); a pest of *Gleditsia: Dasineura gleditchiae* (2011), a pest of *Buxus: Cydalima perspectalis* (2012); a pest of *Albizia: Acizzia jamatonica* (2014); a pest of *Cercis: Cacopsylla pulchella* (2014). Probably most of insects were introduced with imported planting material during the landscaping of the city of Sochi in preparation for the Olympic Games (held in 2014). Quarantine measures should be taken to prevent dispersal of these pests to other regions of the Caucasus and countries of the Black Sea region. Attention should be paid to a new pest for Europe *Medythia nigrobilineata*.

## Introduction

The rapid spread of invasive pests is a great economic and ecological problem of the 21^st^ century (Beenen and Roques 2010). In 2011-2017 an unusually high number of invasive pests new to European Russia were firstly detected in subtropics of the Northwest Caucasus, namely in the city of Sochi.

We present the first reports of two pests new for the Caucasus and European Russia found in 2017: *Drosophila suzukii* and *Otiorhynchus armadillo* and review of other 18 invasive pests firstly recorded for Russia in Sochi in the last 7 years. Most of this information has not been included in the EPPO database (2017) and was published in Russian only.

## Materials and methods

The collecting site, the city of Sochi, is located in the south of European Russia near the Black Sea Coast. This large Russian holiday resort occupies a long narrow band (145 km) between the mountains and the sea. The climate is humid subtropical with a warm rainy winter and a sunny summer. The average annual temperature is +13.6 ^°^C. The average annual precipitation is 1703 mm. The coldest month in the city is February with an average temperature of +6.0°C. The warmest month is August, its average daily temperature is +23.0 °C (Mosiyash and Lugavtsov 1967). Twenty two traps made of plastic bottles and baited with a mixture of commercially available red wine and vinegar were placed in different parts of Sochi and kept from 4 to 19 of June 2017. This bait for *Drosophila suzukii* was recommended by (Cini et al. 2012). One more trap baited with grapes harvested by us in Sochi was kept from 18 to 30 of September 2017. *Otiorhynchus armadillo* was collected by hand.

## First reports of invasive pests found in 2017

### *Drosophila suzukii* (Matsumura, 1931) (Diptera, Drosophilidae)

Spotted-wing *Drosophila* is one of the most important invasive pests of fruit-production and wine-production (Cini et al. 2012). A special issue of the Journal of Pest Science was devoted to this species (Biondi et al. 2016). Unlike other vinegar flies, it is able to oviposit and develop in healthy ripening soft fruits. This species is native to Eastern Asia and has recently spread to Europe and the Americas. Since 2011 *D. suzukii* has been included in the EPPO A2 List (EPPO 2017). It has been recorded in at least 20 European countries. In 2014 *D. suzukii* was firstly recorded for the Ukraine on the Black Sea coast of Crimea (Lavrinienko et al. 2017).

We present the first record of this pest in European Russia and the Caucasus. One male *D. suzukii* was found in the trap kept near the railway station Sochi (43°35'N, 39°44'E) from 4 to 19 of June 2017. One mature female and one female from puparium was found in the trap kept in Serafimovicha Street (43°34'N, 39°45'E) from 18 to 30 of September 2017. The species was identified by referring to Hauser (2011) and checking the following characters: male: one black spot at the apical part of wings and two black combs at the apex of 1st and 2nd tarsal segments (one comb on each segment), female: ovipositor with many dark sclerotized teeth; ovipositor is much longer (at least 6X) than spermatheca diameter.

Since the specimens were captured in different localities and in different months, it indicates, that a breeding population exists. Wine production and fruit production are important branches of economy of the Caucasus region. So the establishment of *D. suzukii* could cause serious negative consequences.

### *Otiorhynchus armadillo* (Rossi, 1792) (Coleoptera, Curculionidae)

We were informed that extensive ornamental plantations of *Viburnum tinus* in Imeretian Resort (Sochi, 43°25'N, 39°56'E) had been severely damaged by an unknown pest. During a survey on 8 June 2017 we found that many leaves had feeding damage that was characteristic for Curculionidae (semicircular cuts on edges) and we collected one female of *Otiorhynchus armadillo.* Similar damage has been noted also on *V. tinus, Viburnum rhytidophyllum, Photinia fraseri* and *Osmanthus heterophyllus* in 2015-2017. Seedlings of *Viburnum tinus* and other damaged plants were imported from Italy in 2012, the pest was obviously introduced with them.

The specimen was identified by referring to Heijerman and Hellingman (2008). The distinctive features are: body black; dorsum covered by yellowish hairs, which arranged in small patches on elytra; rostrum with one high medial keel, without medial furrow; antennae thin, 1st and 2nd funicular antennomeres very thin, elongate, antennomeres 3 to 7 distinctly longer than wide; pronotum covered by dense granules; elytra broadly oval, greatly expanded from the base, slightly narrowed to the apex; shoulders more developed than in *O. salicicola;* intervals covered by irregular granules partly fused to each other and forming transverse wrinkles; striae with more or less regular rows of granules, 1st and 2nd striae connected with each other at elytral apex, all intervals evenly convex, without keels in odd intervals, fore tibiae without external apical lobe. Our specimen is 9.8 mm long.

*Otiorhynchus armadillo* is a pest of trees and shrubs including *Viburnum.* Adults feed on the leaves of 25 families of plants, the larvae feed on roots (Heijerman and Hellingman 2008). Before the 1990s the distribution was restricted to Italy, France, Germany, Croatia, Austria and Belgium (Mazur and Mokrzycki 2011). This flightless weevil has now also been introduced with seedlings to the UK, the Netherlands, Poland, Norway, Czech Republic, Slovakia and Turkey. It is also known from Hungary, Romania, Slovenia, Sweden, Greece, Liechtenstein, Switzerland (Magnano and Alonso-Zarazaga, 2013; Çerçi 2016). In the Netherlands and the UK it has become a serious pest (Heijerman and Hellingman 2008; Halstead 2011).

## Invasive pests found in 2011-2016

*Halyomorpha halys* Stål, 1855 (Hemiptera, Pentatomidae) is a very damaging invasive polyphagous pest native to East Asia and established in North America and Europe in the 1990s and 2000s, respectively (Haye and Weber 2017). In 2008-2013 *H. halys* was included in EPPO Alert List, and since 2002 it has been included in the NAPPO Alert List (EPPO 2017). It was firstly found in Russia in 2014 in Sochi. Since 2015 a severe outbreak is being observed, which has led to significant losses of harvest of fruits (Gapon 2016; Karpun et al. 2017). Recently this pest was recorded in different parts of Krasnodar Territory, Adygea and Abkhazia.

*Paysandisia archon* (Burmeister, 1880) (Lepidoptera, Castniidae) is native to South America. This serious invasive pest of palm trees is included in A2 EPPO list and NAPPO Alert List. It was firstly recorded in Europe (France) in the 1990s and then spread to Belgium, Cyprus, Greece, Italy, Spain and Switzerland (EPPO 2017). It was firstly recorded in Russia in Sochi in 2014 on *Trachycarpus fortunei* and in 2015 spread all over the city and killed a lot of trees: *T. fortunei, Washingtonia filifera, W. robusta, Phoenix canariensis* and *Chamaerops humilis* (Karpun et al. 2015b).

*Rhynchophorus ferrugineus* (Olivier, 1791) (Coleoptera, Dryophthoridae) is native to South-East Asia. In the 1980s this began to spread outside its native range and by now has established in the Middle East, North Africa, North America, countries of the Caribbean and South Europe (EPPO 2017). It was added to the EPPO A2 list in 2006 and is regarded as a quarantine pest in several countries (EPPO 2017). In 2009 *R. ferrugineus* was found in the Caucasus for the first time in Georgia (Pelikh 2009). In 2012-2013 it was recorded in Sochi, where it mainly damages *Phoenix canariensis* and *Washingtonia robusta* (Karpun et al. 2014).

*Epitrix hirtipennis* (Melsheimer, 1847) (Coleoptera, Chrysomelidae) is a pest of tobacco and other plants of the family Solanaceae. It is native to the Americas and established in Europe in the 1980s. It was first recorded for Russia and the Caucasus in Sochi and Tuapse in 2013 (Orlova-Bienkowskaja 2014) and then repeatedly found in Sochi in 2016 (our unpublished data). This pest could be introduced to the Caucasus with seedlings, because its larvae develop on roots. There is no information about the economic importance of this species in the region.

*Dinoderus japonicus* Lesné, 1895 (Coleoptera Bostrichidae) is a serious pest of bamboo native to East Asia. It was often intercepted in Europe with imported bamboo products, but did not establish until recently. In 2012-2013 established populations were found in France and Italy (Nardi et al. 2015). In 2016 it was recorded in European Russia and the Caucasus for the first time: an established population was found in Sochi in the in the thicket of *Phyllostachys* sp. (Bieńkowski and Orlova-Bienkowskaja 2017).

*Luperomorpha xanthodera* (Fairmaire, 1888) (Coleoptera, Chrysomelidae) is a pest of ornamental flowers. Adults feed on flowers, whereas larvae develop on roots. The species is native to China and Korea, established in Europe in 2003, and since then quickly expanded its range (Beenen and Roques 2010). It was recorded for the first time in the Caucasus and Russia in 2016 and severely damages roses and other flowers in Sochi (Bienkowski and Orlova-Bienkowskaja 2017). Larvae could probably be introduced to the region with seedlings.

*Medythia nigrobilineata* (Motschulsky, 1861) (Coleoptera, Chrysomelidae) is serious pest of soybeans in Asia. It is native to China, Far East of Russia, Japan, Nepal, Pakistan and South Korea and had not yet been recorded in Europe before 2016 (Beenen 2010; Toepfer et al. 2014). In 2016 a single female of *M. nigrobilineata* was collected on ruderal plants in Sochi (Bieńkowski and Orlova-Bienkowskaja 2017).

*Harmonía axyridis* (Pallas, 1773) (Coleoptera, Coccinelidae) is native to East Asia. It has been used for biological control of aphids all over the world and has become almost cosmopolitan (EPPO 2017). In some regions *H. axyridis* is regarded as a pest of wine production and fruit production, since it can feed on fruits. An established population was first recorded in Sochi in 2012 (Belyakova and Reznik 2013).The species is now common in Sochi (Ukrainsky and Orlova-Bienkowskaja 2014 Orlova-Bienkowskaja 2014), but it is unknown, if it has an economic impact.

*Stelidota geminata* (Say, 1825) (Coleoptera, Nitidulidae) is a pest of strawberry in its native range, in North America (Connell 1980). Since the 1980s it has been spreading in Europe, though only a few cases of damage of strawberry have been recorded (Spasić et al. 2011). In 2013 the species was found for the first time in the Caucasus in Sochi and Abkhazia (Tsinkevich and Solodovnikov 2014). It now is common in Sochi on decaying fruits.

*Ophelimus maskelli* (Ashmead, 1900) (Hymenoptera, Eulophidae) damaging leaves of many *Eucalyptus* species, began to spread outside its native range (Australia) at the beginning of the 2000s. It has spread to North America, North Africa, the Middle East, and South Europe (EPPO 2017). The pest was first recorded in Russia and the Caucasus in Sochi in 2011 and became abundant in 2013 (Karpun et al. 2015a). Now it is common in Sochi and damages *Eucalyptus cinerea, E. gunii, E. globulus* and *E. viminalis.*

*Glycaspis brimblecombei* Moore, 1964 (Hemiptera, Psyllidae) is another Australian pest of *Eucalyptus.* In 1998 it was firstly recorded outside its native range (in North America) and then appeared also in South America, South and North Africa, Mediterranean countries and South Europe. *Glycaspis brimblecombei* was included in the EPPO Alert List between 2002 and 2006 (EPPO 2017). In 2014 it was first recorded in Russia in Sochi, where it damages leaves of *Eucalyptus viminalis* and *E. globulus* (Karpun et al. 2015a).

*Leptocybe invasa* Fisher & La Salle, 2004 (Hymenoptera, Eulophidae) is an invasive gall inducer on *Eucalyptus.* It is supposed that the species is native to Australia, since it is a specialized pest of a plant native to Australia, but it has not yet been found in this continent (EPPO 2017). It was discovered in 2000 in the Middle East and then recorded in the Americas, Africa, Asia and South Europe. The pest was included in EPPO Alert List between 2006 and 2010. In 2014 *L. invasa* was recorded in Sochi (Karpun et al. 2015a). It severely damages *Eucalyptus viminalis* and close species.

*Lamprodila festiva* (Linnaeus, 1767) (Coleoptera, Buprestidae) is a serious pest of ornamental Cupressaceae native to Mediterranean region and South Europe and has recently expanded its range in Europe. After the first record in Sochi in 2013 the species has become common in the region and severely damages *Thuja, Chamaecypaeis, Juniperus* and *Cupressus* (Volkovitsh and Karpun 2017).

*Cydalima perspectalis* (Walker, 1859) (Lepidoptera, Crambidae) is a pest of boxwood native to Asia. It was first found in Europe in 2006 and then spread to 20 countries (EPPO 2017). Between 2007 and 2011 it was included into the EPPO Alert List. *Cydalima perspectalis* was first intercepted in Russia in 2012, and in 2013 became abundant pest all over the Black Sea coast from the border of Abkhazia to Tuapse (Karpun et al. 2015a) as well as in city parks of the Absheron District of Krasnodar Region and the city of Grozny (Chechnya). In 2014 it was recorded in Abkhazia and Georgia, in 2015 also in Crimea and Adygea. It has completely destroyed *Buxus sempervirens* and other boxwood species in Sochi as well as natural plantations of an endemic protected species *B. colchica* (Karpun et al. 2015a).

*Acizzia jamatonica* (Kuwayama, 1908) (Hemiptera, Psyllidae) is a pest of *Albizia* spp. In the 1980s the pest native to Japan began to spread to other countries of Asia. After an accidental introduction to Italy in 2001 it spread to 13 European countries. Since 2006 *A. jamatonica* has been expanding its range in North America. It was included on the EPPO Alert List between 2004 and 2006 (EPPO 2017). In 2014 it was found in Sochi and severely damaged *Albizia* trees (Karpun et al. 2015a).

*Cacopsylla pulchella* (Löw, 1877) (Hemiptera, Psyllidae) is a species of Mediterranean origin, a pest of *Cercis.* In the second half of the twentieth century *C. pulchella* spread widely across Europe (Halperin et al. 1982). It was first recorded in Russia in 2014 in Sochi (Karpun et al. 2015a) and in 2016-2017 severely damaged *Cercis* in city parks.

*Ceroplastes ceriferus* (Fabricius, 1798) (Hemiptera, Pseudococcidae) is a polyphagous pest of trees and shrubs. It is native to India and has become almost cosmopolitan (EPPO 2017). In Europe it was established only in Italy. Between 2002 and 2005 the species was included in the EPPO Alert List. In 2015 *C. ceriferus* was first recorded in Russia in Sochi and became abundant on many plants (Karpun et al. 2017).

*Glyphodes pyloalis* Walker, 1859 (Lepidoptera, Crambidae) *(Daiphania pyloalis)* is a pest of *Morus* native to the USA and Mexico. It is a serious invasive pest of silk-production in Asia. In the 2000s it appeared in Europe in Moldova, Romania, Georgia, Azerbaijan and became an economically significant pest (EPPO 2017). In 2015 *G. pyloalis* was recorded in European Russia for the first time in Sochi (Karpun et al. 2017).

## Discussion

Plantations of Sochi consist mainly of exotic plants. Alien insects are often introduced there with imported planting material, and the subtropical climate promotes their establishment (Karpun et al. 2015a). The rate of invasions has never been as high as now. Previously 5-6 new alien pests per decade appeared. Between 2011 and 2017 at least 20 new invasive species were recorded.

Insects that have recently appeared in the region are native to different continents: Asia (10 species), Australia (3 species), Europe (3 species), North America (3 species) and South America (1 species). Many of them began to spread outside their native ranges 10-20 years ago and spread in different continents or even all over the world. All species except *Medythia nigrobilineata* occur in other European countries. Most of the invasions were probably connected with an import of planting material from European nurseries during the landscaping the city of Sochi in preparation for the Olympic Games in 2011-2013. The exporting countries were mainly Italy, Spain and Montenegro (Karpun et al. 2017). *Drosophila suzukii* and *Halyomorpha halys* could have been unintentionally introduced with imported fruits, *Dinoderus japonicus -* with bamboo products, *Medythia nigrobilineata -* with soybeans. *Glyphodes pyloalis* and *Harmonia axyridis* could also be introduced or spread spontaneo9usly from neighboring countries.

The information about first records of the quarantine pests *Drosophila suzukii, Paysandisia archon* and *Rhynchophorus ferrugineus* in Russia should be included in the EPPO database as well as information about records of other invasive pests. Quarantine measures should be taken to prevent dispersal of pests to other regions of the Caucasus and countries of the Black Sea region. Special attention should be payed to invasion of *Medythia nigrobilineata* to the Caucasus, since this pest is new to Europe.

## Acknowledgements

We are grateful to L.Yu. Ivanova, S.O. Galkina and A.A. Galkin for the help and to D. Eyre (Defra, UK) for linguistic corrections. The study was supported by Russian Science Foundation, Project No 16-14-10031.

